# The S1^B^-Targeting PEDV Neutralizing Antibody C62 Prematurely Activates S-trimer and Mimics Receptor Function

**DOI:** 10.64898/2026.08.05.742767

**Authors:** Jianbo Liu, Sheng Wang, Zimu Li, Mengzhen Su, Zhichen Wang, Ying Li, Jiahui Zou, Xuerui Zhu, Heying Li, Yong Ma, Mei Li, Min Zhou, Yutong Liu, Jingjing Wang, Jun He, Chaofeng Yang, Shaobo Xiao, Xinwen Chen, Hongbo Zhou, Xiaoli Xiong

## Abstract

Porcine epidemic diarrhea virus (PEDV) causes devastating enteric disease in piglets, yet the mechanistic basis of antibody-mediated neutralization remains poorly understood. Here, we determined the structure of PEDV HNXX-strain spike domain B (S1^B^) simultaneously bound by C62, a neutralizing porcine monoclonal antibody against PEDV G2 strains, and N34, a non-neutralizing porcine PEDV antibody. The structure reveals that C62 targets a conserved, cryptic epitope that is accessible only when S1^B^ adopts an “up” conformation. Functionally, we showed that C62 has substantially stronger activity than N34 in triggering S-trimer disassembly and inducing the formation of proteinase K-resistant, post-fusion-like S2 structures. Despite the weaker triggering activity of N34, both C62 and N34 can function as artificial receptors. Notably, although the C62 epitope is conserved across both G1 and G2 strains, C62 exhibits G2-strain-biased neutralizing activity. We further showed that differences in cell-surface membrane fusion activity among PEDV spikes correlate with distinct viral entry pathways and are jointly determined by the S1^A^ and S1^B^ sequences. Together, our findings identify a strain-specific vulnerable site on the PEDV S-trimer and provide insight into how cell-surface membrane fusion activity may influence viral entry pathway selection and antibody neutralization efficacy.

## Introduction

Porcine epidemic diarrhea virus (PEDV), an enteric alphacoronavirus that primarily infects piglets, is a globally prevalent pathogen that causes severe morbidity and mortality in swine, resulting in substantial economic losses in the swine industry^1^. First identified in the United Kingdom in 1971 and subsequently isolated as the CV777 strain in 1978^2,3^, PEDV has circulated in swine populations for over five decades.

Prior to 2010, PEDV strains were genetically homogeneous, classified as genotype 1 (G1) with CV777 as the prototype. However, a novel genotype (G2) emerged in 2010 and rapidly spread, including to the United States^4,5^. G2 strains are characterized by insertions and deletions (INDELs) in the spike (S) gene^4^. Recent genomic analyses reveal further diversification, with S gene lengths now ranging from 4,125 to 4,182 nucleotides [except for strains TC-PC177 (KY499261)^6^ and CN19 (OM814174), which lack 591 and 177 nucleotides, respectively] due to in-frame deletions or insertions of tri-nucleotide motifs, leading to the classification of circulating strains into nine distinct groups^7^.

The S-protein of PEDV is a surface glycoprotein essential for viral entry^8^. As the primary target of neutralizing antibodies induced by infection or vaccination^9,10^, the S-protein is a key focus for subunit vaccine development. Structurally, the PEDV S-protein is synthesized as a single polypeptide and can be divided into the receptor-binding S1 subunit and the membrane-fusion S2 subunit, but unlike sarbecovirus^11^ and merbecovirus^12^ spikes, it lacks a defined S1/S2 cleavage site^11–13^. Toward the C-terminus of the S2 subunit, a transmembrane domain (TM) anchors the S-trimer to the viral envelope, followed by a small cytosolic tail^14^. The S1 subunit consists of five domains: domain 0 (D0 or S1^0^), S1^A^ (also named N-terminal domain, NTD), S1^B^ [corresponds to the receptor binding domain (RBD) in SARS-CoV-2 S-trimer, also named C-terminal domain, CTD], S1^C^ (also named domain C or subdomain 1, SD-1) and S1^D^ (also named domain D or subdomain 2, SD-1)^8,14,15^. Notably, the D0 is a structural feature present in the S-trimers of many alphacoronaviruses but is absent in other coronavirus genera^16^. This domain is known to adopt dynamic “up” (D0_U_) and “down” (D0_D_) conformations, which have also been described in the alphacoronavirus CCoV-HuPn-2018 as “swung-out” and “proximal” positions relative to the S-trimer core, respectively^8,17^. In PEDV, the S1^B^ can likewise transition between “closed” and “open” conformations^8^. The combined dynamics of D0 and S1^B^ generate multiple prefusion S-trimer states. When S1^B^ is closed, the S-trimer can adopt a spectrum of D0 arrangements, including three D0-down protomers (D0_DDD_), two D0-down and one D0-up protomer (D0_DDU_), one D0-down and two D0-up protomers (D0_DUU_), or three D0-up protomers (D0_UUU_). In addition, coupling of these variable D0 arrangements with either open or closed S1^B^ states further expands the conformational landscape of the PEDV S-trimer^8^.

To date, most PEDV-neutralizing antibodies have been reported to recognize epitopes within the S1 subunit, and only a single S2-specific antibody has been described^10,18–21^. However, the molecular mechanism underlying their neutralization activities remain incompletely understood. Because PEDV is now divided into two distinct genotypes, G1 and G2^5^, the observed spectrum of antibody breadth, ranging from genotype-restricted to pan-genotypic activity, suggests that neutralization may be governed by distinct epitope-dependent mechanisms. Thus, a comprehensive mechanistic understanding of PEDV antibody neutralization is still lacking.

Previous studies have shown that the antibody C62 binds within the S1^B^ of the PEDV S-protein and potently inhibits infection by multiple G2 strains^10^. To elucidate the mechanistic basis of C62-mediated neutralization, we determined a cryo-electron microscopy (cryo-EM) structure of PEDV S1^B^ bound to C62, with the non-neutralizing antibody N34 also bound and used as a molecular-weight fiducial. Integrated with functional analyses, this structure reveals how C62 blocks viral entry and highlights an S1^B^ dynamics-dependent mechanism that influences both antibody neutralization and viral entry pathway usage.

## Results

### C62 effectively neutralizes PEDV G2 strains

In our previous work, we isolated a broadly neutralizing porcine monoclonal antibody, C62, using single B cell sequencing technology^10^. This antibody was shown to effectively neutralize multiple PEDV G2 strains, including HNAY, WHLL, HNXX, ECQ1, EHUB4, AJ1102, and YN150, which distribute in several different subgroups of G2 (**Fig. S1**)^10^. However, C62 neutralization of the classical G1 strain CV777 had not been evaluated. We therefore tested C62 against CV777 alongside the G2 strain HNXX. C62 potently neutralized HNXX (IC_50_ = 15.8 μg/ml) but only weakly neutralized CV777 (IC_50_ = 127.5 μg/ml) (**Fig. 1A and 1C**), confirming its G2-biased neutralization profile and highlighting its broad activity against diverse G2 variants.

**Fig. 1.**
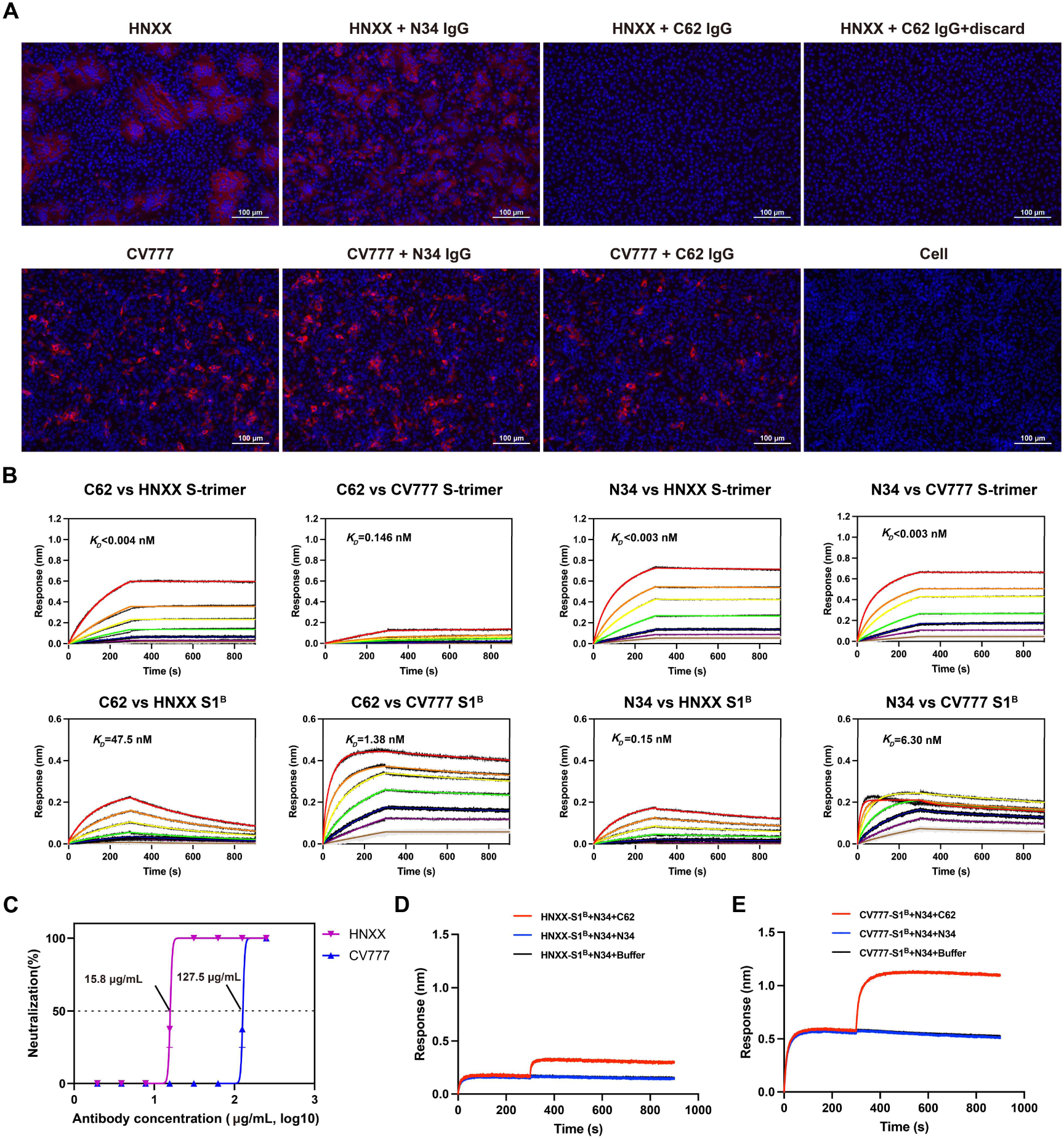
Characteristics of S1^B^ targeting PEDV mAbs C62 and N34. **A**, Neutralization potencies of C62 and N34 against PEDV strains CV777 and HNXX in cell culture. For mAb C62, a condition in which the antibody was discarded after the initial incubation period was also tested (HNXX + C62 IgG + discard). Scale bars, 100 μm. **B**, BLI sensorgrams of C62 and N34 IgG binding to HNXX and CV777 S-trimers or S1^B^ proteins. Two-fold serial dilutions of S-trimer or S1^B^ (200 nM to 3.125 nM) were used for kinetic measurements. Detailed kinetic parameters are provided in **Table S1. C,** Neutralization curves of C62 against PEDV strains HNXX and CV777 with IC_50_ values shown. **D**, Competition binding assay between C62 and N34 for HNXX S1^B^. **E**, Competition binding assay between C62 and N34 for CV777 S1^B^.

Previously, we established that C62 blocks viral attachment, as evidenced by qPCR-based assays in which persistent presence of the antibody in culture medium prevented virus infection^10^. Here, we further explored its mechanism. After Vero cells were co-incubated with HNXX virus-C62 mixture for 1 h, the inoculum was replaced with antibody-free medium. Under these conditions, no PEDV-positive cells were detected by staining with an N-protein-specific antibody (**Fig. 1A**). In contrast, the control antibody N34, which also binds PEDV S1^B^ but lacks neutralizing activity, permitted widespread infection. These results indicate that C62 neutralizes viral infection by blocking viral attachment and/or membrane fusion, thereby acting at an early stage of viral entry.

### C62 binds both G1 and G2 PEDV S-proteins

Using bio-layer interferometry (BLI) assay, we immobilized C62 on the biosensor and tested binding of C62 to the S1^B^ monomer and S-trimer of the HNXX strain. C62 binds S1^B^ monomer with an affinity of 48 nM. Binding to the HNXX S-trimer was greatly amplified likely by avidity^22^, showing largely non-dissociating binding curves, with an affinity estimated to be tighter than 4.0 pM. Interestingly, we were surprised to find that C62 binds CV777 S1^B^ monomer with even greater affinity of 1.4 nM. However, binding to CV777 S-trimer is significantly weaker, showing binding curves of greatly reduced responses and an affinity of 0.15 nM (**Fig. 1B**), which may explain why C62 fails to neutralize this strain. We also tested binding of the non-neutralizing antibody N34 targeting S1^B^. N34 is able to bind both HNXX and CV777 S1^B^ monomer and S-trimers. We also show that C62 and N34 bind HNXX and CV777 S1^B^ monomer non-competitively (**Fig. 1D and 1E**).

### Structural basis of C62 binding to PEDV S1^B^

Although purified HNXX PEDV S-trimer produced in Expi293F cells bound strongly to C62 in BLI assays (**Fig. 1B**), cryo-EM analysis of the HNXX S-trimer:C62 Fab sample failed to identify the expected S-trimer:Fab complexes; instead, the sample mainly contained unbound S-trimers and disintegrated particles, making it unsuitable for structure determination.

Since C62 binds the S1^B^ of the S-trimer with high affinity (**Fig.1B**), we next attempted to form a complex between the S1^B^ and the C62 Fab. To facilitate structural determination, the non-competing N34 Fab was further included to increase the molecular weight of the complex as a fiducial (**Fig.1D**), following previously used strategies^23–25^.

Cryo-EM analysis of the C62 Fab:S1^B^:N34 Fab ternary complex yielded a reconstruction at an overall resolution of 3.0 Å resolution (**Fig. S2 and S3**), clearly resolving binding interfaces between S1^B^ and both Fabs. This structure shows that C62 recognizes a conformational epitope on S1^B^ (**Fig. 2E and F, Fig. S3 and S4**), burying a total surface area of 679.3 Å^2^, with 525.8 Å^2^² contributed by heavy-chain complementarity-determining regions (CDRs) (HCDRs) and 153.5 Å^2^ by light-chain CDRs (LCDRs; **Fig. 2A to D**). The majority of contacts are mediated by HCDR1–3 and LCDR2 (**Fig. 2E to G**).

**Fig. 2.**
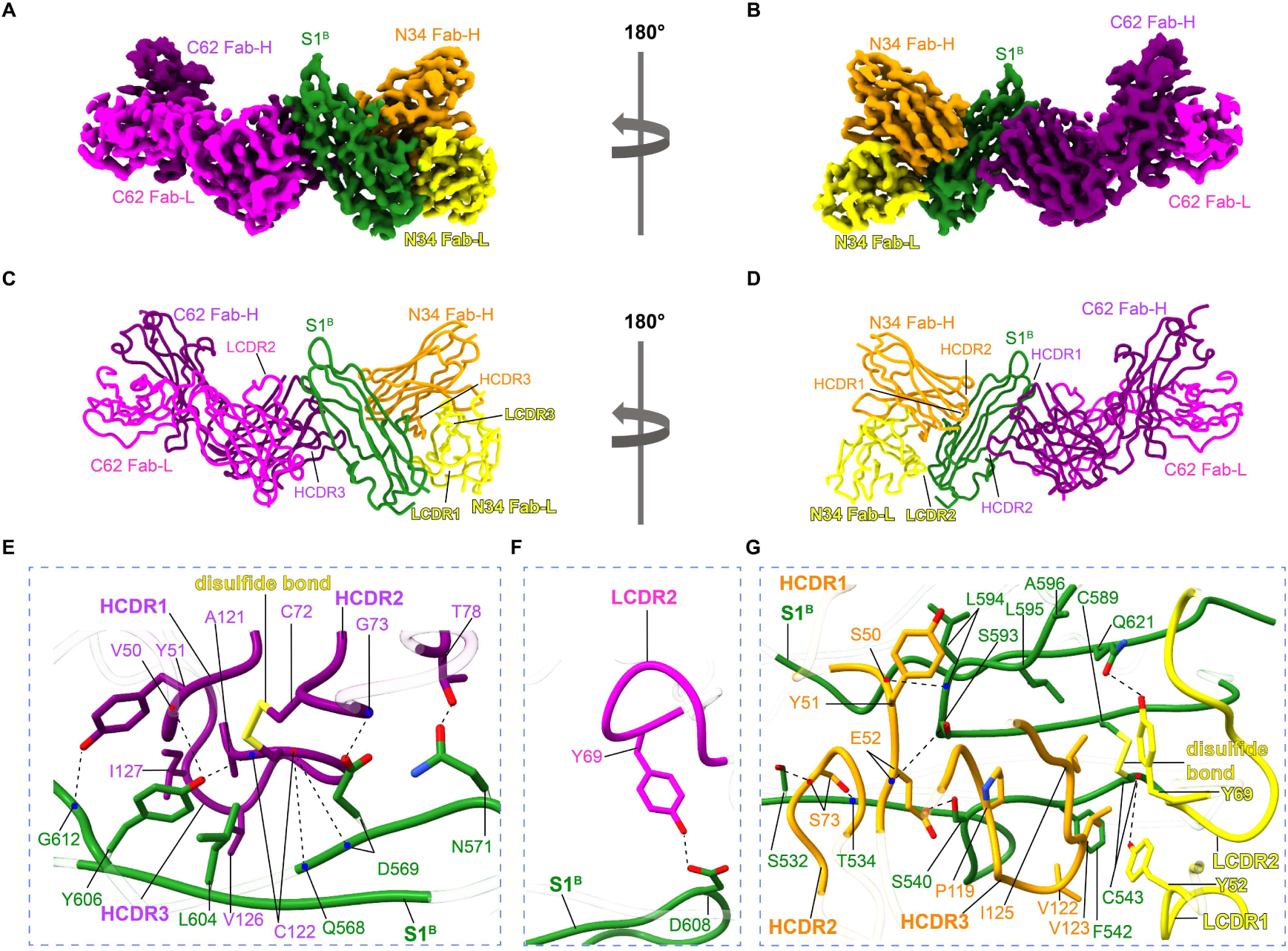
Structural basis of PEDV S1^B^ recognition by C62 and N34 Fabs. **A** and **B**, Electron density map of the C62 Fab:S1^B^:N34 Fab ternary complex. The map is shown in two orientations, rotated 180° relative to each other, to visualize antibody binding. Protein chains are colored as follows: C62 Fab heavy chain (Fab-H) (purple), light chain (Fab-L) (magenta); N34 Fab-H (orange), Fab-L (yellow); S1^B^ (dark green). **C** and **D**, The corresponding C62 Fab:S1^B^:N34 Fab ternary complex molecular model in the same orientations of **A** and **B**. **E** and **F**, Detailed views of the binding interfaces between C62 Fab and S1^B^. Hydrogen bonds are shown in dotted line. **G**, Detailed view of the binding interface between N34 Fab and S1^B^.

Notably, mAb C62 utilizes an unusual inter-CDR disulfide bridge between C72^H^ (HCDR2) and C122^H^ (HCDR3) to stabilize a large exposed hydrophobic surface to bind S1^B^ residues Y606 and L604. This hydrophobic surface is formed by residues from HCDR1 (V50^H^, and Y51^H^), HCDR2 (C72^H^, and G73^H^), and HCDR3 (A121^H^, C122^H^, G123^H^, G124^H^, G125^H^, V126^H^, and I127^H^). Reinforcing this hydrophobic contact is a sophisticated hydrogen bond network. The mainchain carbonyl of V50^H^ (HCDR1) and the mainchain amide of C122^H^ (HCDR3) each forms a hydrogen bond with the sidechain hydroxyl of Y606 (S1^B^). The mainchain carbonyl of HCDR3’s C122^H^ hydrogen-bonds with the backbone amides of Q568 and D569 in S1^B^; further, the C62 G73^H^ mainchain amide interacts with D569 sidechain, while T78^H^ sidechain forms a hydrogen bond with S1^B^ residue N571. On the light chain, the LCDR2 residue Y69^L^ form a hydrogen bond with D608 of S1^B^ (**Fig. 2E to F**).

### Structural basis of N34 binding to PEDV S1^B^

Although N34 does not neutralize PEDV, it also recognizes a conformational epitope on the surface of S1^B^ opposite the C62 epitope (**Fig. 2G, Fig. S3 and S4**). N34 buries a total surface area of 1009.9 Å^2^. Of which, 653.6 Å^2^ is contributed by HCDRs and 356.3 Å^2^ by LCDRs (**Fig. 2A to 2D**).

Notably, mAb N34 primarily utilizes two distinct hydrophobic surfaces to engage S1^B^. The first surface is formed by LCDR1 (Y52^L^), LCDR2 (Y69^L^), and HCDR3 residues (P119^H^, V122^H^, V123^H^, and I125^H^). This surface engages S1^B^ residues F542, C543, C589, L595, and A596, including the disulfide bond between C542 and C589. The second surface is formed by Y51^H^ (HCDR1) and engages S1^B^ residue L594. These hydrophobic contacts are further stabilized by an extensive hydrogen bond network. Specifically, the mainchain carbonyl of S50^H^ in HCDR1 forms a hydrogen bond with the mainchain amide of L594 in S1^B^. E52^H^ (HCDR1) mainchain amide forms a hydrogen bond with the S593 sidechain while E52^H^ sidechain forms a hydrogen bond with S540 of S1^B^. S73^H^ (HCDR2) forms hydrogen bonds with S532 and T534. LCDRs also contribute: the sidechain hydroxyl of Y52^L^ (LCDR1) hydrogen-bonds with the C543 mainchain carbonyl, while Y69^L^ (LCDR2) forms a hydrogen bond with Q621.

### S-trimer conformation dictates the accessibility of the C62 and N34 epitopes

In the PEDV S-trimer, the S1^B^ has been shown to adopt both “down” (or “closed”) and “up” (or “open”) conformations when D0 is in the “up” conformation^8^. Structural analysis revealed that, because C62 approaches S1^B^ from below, its epitope is fully buried when S1^B^ adopts the “down” conformation but becomes exposed upon transition to the “up” conformation (**Fig. 3A, and D**). Docking of the C62 Fab onto the “3-S1^B^-down” S-trimer conformation revealed steric clashes with S1^A^ and the neighboring S1^B^. In contrast, docking C62 Fab onto the “1-S1^B^-up” and “3-S1^B^-up” S-trimer showed no apparent clashes (**Fig. 3C, E to G**). Therefore, C62 requires the S1^B^ to adopt an “up” conformation before the antibody can bind (**Fig. 3H to J**).

**Fig. 3.**
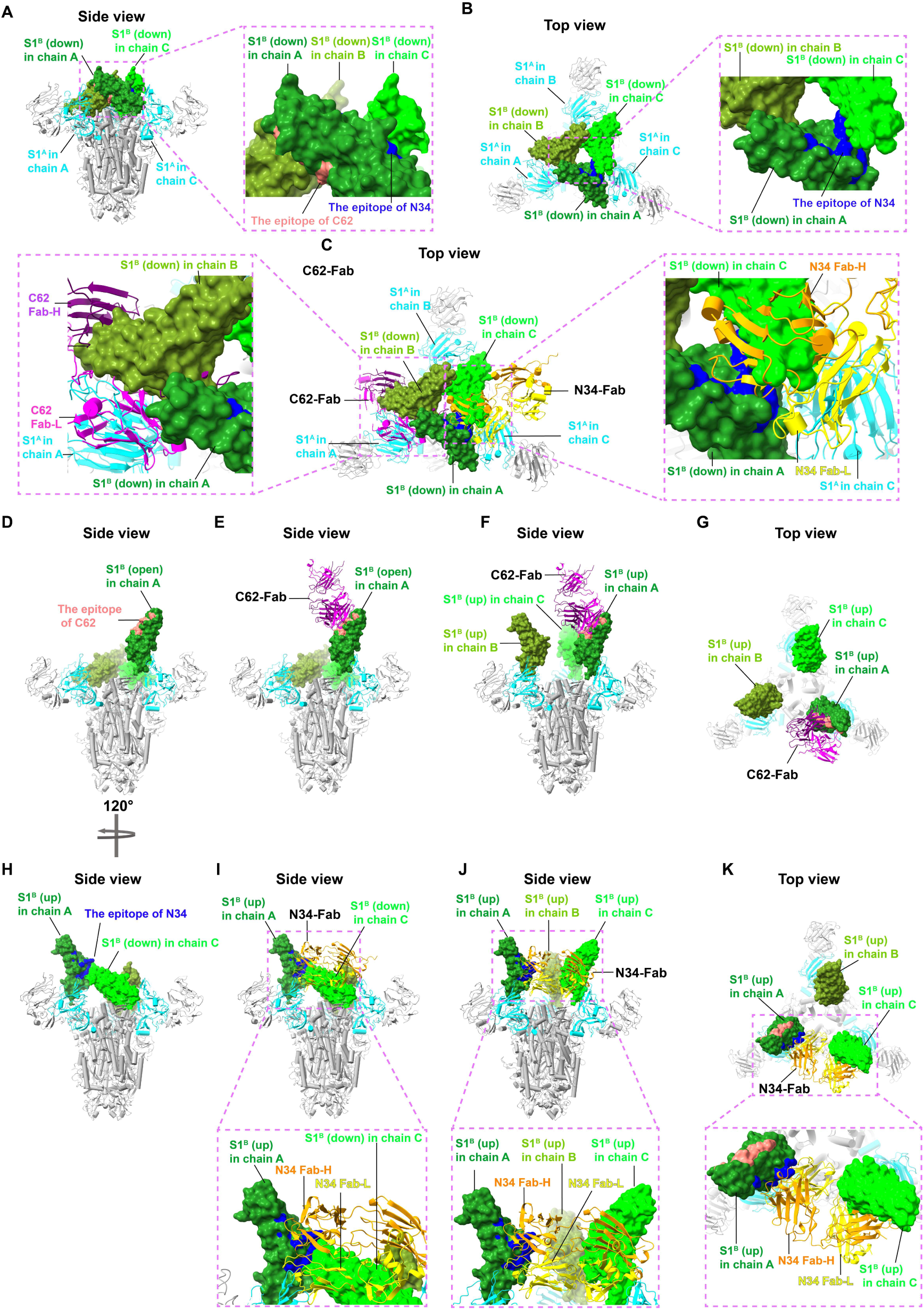
Steric hindrance analysis of C62 and N34 Fab binding to the PEDV S-trimer. S-trimer structural elements are color-coded as follows: S1^A^ in cyan, S1^B^ in dark green/lime/olive green for the three protomers. The C62 Fab-H/L chains are shown in purple and magenta, respectively, whereas N34 Fab-H/L chains are shown in orange and yellow, respectively. **A** and **B**, Side and top views of the “3-D0-up, 3-S1^B^-down” HNXX S-trimer (PDB: 9XNZ) with epitopes of C62 (salmon) and N34 (blue). Pink dashed boxes in A and B show magnified views of the C62/N34 epitopes. **C**, Top view of the C62 Fab:S1^B^:N34 Fab atomic model docked into the S-trimer (3-D0-up, 3-S1^B^-down conformation; 9XNZ). Pink dashed boxes in C show magnified views of the steric-hindrance regions of C62 (left) and N34 (right). **D, E, H,** and **I**, Side views of an S-trimer with a 3-D0-up, 1-S1^B^-up conformation, modelled using PDB structures 9XNZ and 7Y6V. **D** and **H** highlight the epitopes of C62 (salmon) and N34 (blue), respectively, while **E** and **I** show the corresponding Fab binding modes. **F, G, J,** and **K**, Side views of the S-trimer (3-D0-up) with three S1^B^-open protomers, modelled using PDB structures 9XNZ and 7Y6V. **F** and **G** show side and top views of the C62 Fab binding mode, respectively. **J** and **K** show side and top views of the N34 Fab binding mode, respectively. The pink dashed boxes in **I**–**K** indicate magnified views of the steric hindrance regions in the N34 binding models.

Unlike the C62 epitope, the N34 epitope is semi-exposed when S1^B^ is in the “down” conformation and exposed when in the “up” conformation (**Fig. 3A, B, and H**). In contrast, docking of the N34 Fab onto an “up” S1^B^ within a S-trimer revealed steric clashes with the neighboring S1^B^ regardless of whether it adopted the “up” or “down” position (**Fig. 3C, I–K**). Therefore, although both antibodies demonstrated binding to S-trimers in BLI assays (**Fig. 1B**), structural analysis suggests that the epitopes of both C62 and N34 require S-trimer conformational rearrangements for antibody access, with N34 requiring more extensive changes beyond the S1^B^ domain adopting an “up” conformation.

### C62 depolymerizes and triggers the PEDV S-trimer

Our structural analysis suggests that binding of both C62 and N34 requires conformational changes in the S1^B^. Notably, in SARS-CoV-2 and SARS-CoV-1, certain S1^B^-targeting antibodies can induce S1^B^ (RBD) conformational rearrangements that prematurely trigger the S-trimer, which could contribute to their neutralizing activity^26–28^.

To test whether C62 or N34 could trigger PEDV S-trimer, we incubated purified S-trimers with either C62 or N34. Purified HNXX S-trimer was incubated with antibodies for 1 h at room temperature, followed by trypsin treatment for 1 h at 37 °C. In the absence of antibody, HNXX S-trimer remained largely resistant to trypsin (**Fig. 4A**). Similar resistance was observed in the presence of N34, indicating that the addition of N34 did not markedly alter the protease sensitivity of the S-trimer under the assay conditions (**Fig. 4A**). By contrast, C62 treatment caused the intact S-protein band to disappear almost completely, indicating that C62 induces conformational changes that render the S-trimer susceptible to protease digestion. In addition to the disappearance of the intact S-protein band, co-incubation of the S-trimer with C62 increased the intensity of lower-molecular-weight bands that were resistant to trypsin digestion, likely corresponding to protease-resistant post-fusion S2 structures (**Fig. 4A**). Notably, weak but detectable trypsin-resistant bands were also observed after incubation with N34, appearing slightly more intense than those in the no-antibody control. This suggests that N34 binding may have a limited ability to render the HNXX S-trimer labile to trypsin digestion, although this effect is much weaker than that observed with C62.

**Fig. 4.**
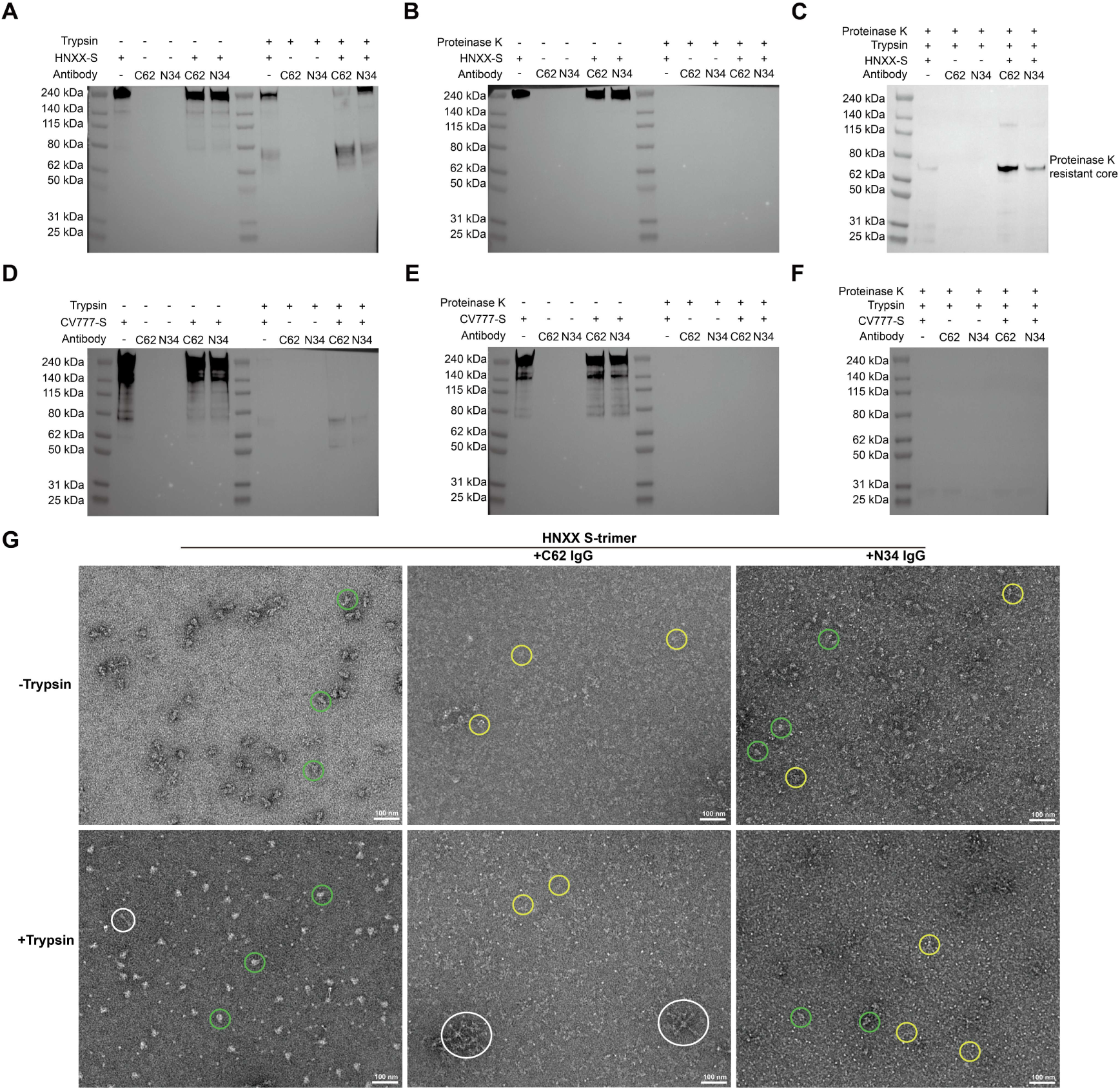
Antibody C62 triggers PEDV S-trimers forming post-fusion like structures. **A and D**, Western blot assessment of C62 IgG-induced perturbation of HNXX and CV777 S-trimer structures in the presence of trypsin. **C and F**, Western blot assessment of C62 IgG-induced perturbation of HNXX and CV777 S-trimer structures following sequential treatment with trypsin and proteinase K. After pre-incubation with either C62 IgG or N34 IgG for 1 h at room temperature, the S-trimer was sequentially digested with trypsin and proteinase K each for 1 h. **G**, Representative negative-stain electron micrographs visualizing the C62-induced disintegration of the HNXX S-trimer. Examples of postfusion structures aggregated into rosettes are outlined in white, S-trimers that have disintegrated into smaller structures are outlined in yellow, and intact S-trimers are outlined in green.

Previous studies have shown that several SARS-CoV-1 and SARS-CoV-2 S1^B^ (RBD)-targeting neutralizing antibodies, including S230, B38, R1-26, and rmAb23, can induce post-fusion S2 structures characterized by resistance to proteinase K digestion^26–28^. Similar proteinase K-resistant post-fusion S2 structures have also been observed for MHV S-proteins^29–31^, suggesting this feature being conserved among coronaviruses. Using western blotting, we confirmed that the trypsin-resistant bands induced by C62 and N34 were also resistant to proteinase K digestion, further supporting their assignment as proteinase K-resistant S2 post-fusion structures (**Fig. 4C**). Consistently, the proteinase K digestion experiment indicates that C62 triggers S-trimer conformational changes much more efficiently than N34. Interestingly, purified CV777 S-trimer was intrinsically susceptible to trypsin digestion under the same conditions used for the HNXX S-trimer, and proteinase K digestion experiments indicate that CV777 S-trimer was only very weakly triggered by C62 and N34 antibodies (**Fig. 4D to F**).

Negative-stain TEM imaging showed that C62 Fab treatment disrupted the HNXX S-trimer structures and promoted the formation of post-fusion-like structures (**Fig. 4G**), providing direct visual evidence for C62-induced premature triggering of the HNXX S-trimer. In contrast, N34 treatment produced fewer such structures, suggesting a weaker ability to trigger S-trimer conformational change (**Fig. 4D**).

### CV777 and HNXX S-proteins mediate distinct viral entry pathways

Our trypsin-digestion experiments suggest that the HNXX S-trimer is largely resistant to trypsin, whereas the CV777 S-trimer is intrinsically trypsin labile. Interestingly, analysis of cells infected with CV777 or HNXX revealed distinct infection-associated morphologies: HNXX induced substantial syncytium formation, whereas CV777 exhibited a markedly reduced syncytium-forming activity compared with HNXX, producing few, if any, syncytium (**Fig. 1A**). These results suggest that surface-expressed CV777 S-trimers mediate much weaker cell-cell fusion than HNXX S-trimers. Therefore, we hypothesized that the two strains may be biased toward distinct cell-entry pathways, as coronaviruses have been generally identified to enter cells either at the plasma membrane or through the endosomal route^32–37^. To test this, we used camostat, a well-characterized serine protease inhibitor that blocks trypsin-like proteases, including TMPRSS2 and trypsin, the latter of which was added during PEDV infection^38^. We also tested E64d, a cysteine protease inhibitor that primary inhibit endosomal cathepsins to block endosomal entry^39^. Camostat markedly reduced HNXX infection to a level comparable to that observed in the no-trypsin DMSO control, suggesting that HNXX requires exogenous trypsin for efficient cell entry and that camostat-mediated inhibition of trypsin strongly impairs HNXX entry. By contrast, the addition of camostat had little effect on CV777 infection. Conversely, E64d substantially reduced CV777 infection but had little effect on HNXX infection (**Fig. 5A**–**5C**). These results suggest that HNXX enters cells primarily through plasma membrane fusion promoted by exogenous trypsin, whereas CV777 relies mainly on the endosomal entry pathway (**Fig. 5A and 5B**).

**Fig. 5.**
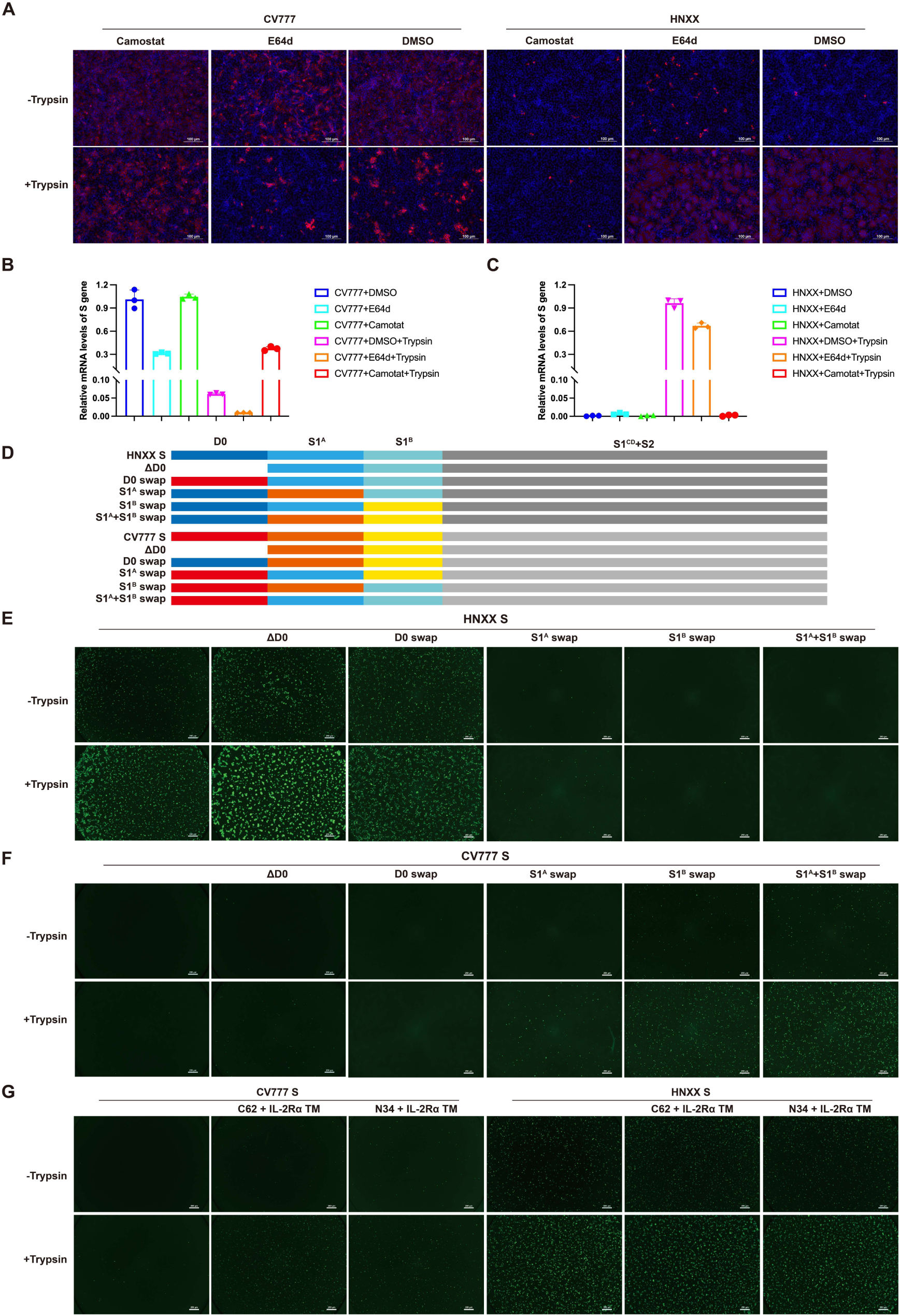
Differential sensitivity of PEDV strains CV777 and HNXX to protease inhibitors and distinct cell-surface fusogenicity of their spike proteins. **A,** CV777 and HNXX strains show different sensitivity to different protease inhibitors. Cells were pretreated with either the serine protease inhibitor camostat or the cysteine protease inhibitor E-64d prior to infection; a DMSO vehicle served as the control. **B and C,** Viral entry efficiency, quantified by qPCR measuring intracellular viral S gene mRNA levels following inhibitor treatment. **D**, Schematic representation of D0-deletion(ΔD0) and domain-swap constructs between CV777 and HNXX S-proteins, involving swapping of the D0, S1^A^, S1^B^, or S1^A^+S1^B^ regions. **E and F**, Cell-cell fusion activity mediated by wild-type or chimeric spike proteins. Constructs included ΔD0 and domain swaps (illustrated in **D**) between the CV777 and HNXX spikes. Assays were performed in the presence or absence of trypsin (1 μg/mL). **G**, the effect of cell-surface expressed C62 or N34 antibody with the IL-2Rα TM fused to the C-terminus of their heavy chains on spike-mediated cell-cell fusion.

To further characterize fusion activities of the two S-proteins, we employed a HEK293T cell-based cell-cell fusion assay^11,12,40^. In this assay, effector cells were co-transfected with PEDV S-protein and the split GFP1-10 fragment, while target cells were transfected with the complementary GFP11 fragment only without overexpressing any receptor. HNXX S-protein mediated weak yet detectable cell-cell fusion in the absence of exogenous trypsin. Cell-cell fusion was greatly amplified after trypsin addition (1 μg/ml) (**Fig. 5E**), suggesting HNXX S-protein mediated efficient cell surface membrane fusion in the presence of exogenous trypsin. The S-protein of CV777 mediated no cell-cell fusion in the absence of trypsin. In the presence of trypsin (1 μg/ml), only very weak fusion was observed, suggesting that CV777 S-protein cannot mediate efficient cell-surface fusion (**Fig. 5F**). These findings are consistent with the differential camostat and E64d inhibition profiles observed for HNXX and CV777 viruses (**Fig. 5A and 5B**). Together, our results indicate that HNXX primarily enters cells through cell-surface membrane fusion, consistent with the cell-surface fusion activity of its spike, whereas CV777 relies mainly on endosomal fusion for entry.

### S1^A^ and S1^B^ cooperatively control PEDV S-trimer mediated membrane fusion

Since the D0 has been reported to restrict S-protein-mediated cell-cell fusion in feline coronavirus (FCoV)^41^, and the D0 exhibits the highest degree of divergence between the two strains, we generated CV777 and HNXX S-proteins with D0-deleted, and employed the cell-cell fusion assay to test their fusion activity. D0 deletion substantially enhanced the fusion activity of the HNXX S-protein, but only marginally increased the fusion activity of the CV777 S-protein (**Fig. 5E and F**). To identify which domain or domains influence the cell-cell fusion, we systematically exchanged D0, S1^A^, and S1^B^ between the two S-proteins and tested the resulting chimeric S-proteins using the cell fusion assay (**Fig. 5D to F, and S5**). Reciprocal domain-exchange analysis revealed distinct contributions of S1 subdomains to S-protein-mediated cell-cell fusion. In the presence of trypsin, which enhances S-protein-mediated fusion, D0 replacement caused only a modest reduction in HNXX S-protein-mediated fusion, whereas replacement of either S1^A^ or S1^B^ markedly impaired fusion. Simultaneous replacement of both S1^A^ and S1^B^ completely abolished HNXX S-protein-mediated fusion. Conversely, for the CV777 S-protein, replacement of D0 had no detectable effect on fusion activity, whereas replacement of S1^A^ mildly enhanced fusion and replacement of S1^B^ led to a marked increase. Notably, simultaneous replacement of both S1^A^ and S1^B^ conferred robust fusion activity over the CV777 S-protein (**Fig. 5E and F**). Together, these data indicate that S1^A^ and S1^B^ cooperatively determine the cell-surface membrane fusion activity in HNXX and CV777 PEDV S-proteins.

### C62 and N34 binding enhances CV777 S-protein-mediated cell-surface fusion activity

The epitopes of C62 and N34 are largely conserved between the HNXX and CV777 S-proteins (**Fig. S4**), and both antibodies bind to the S1^B^ of these two S-proteins (**Fig. 1B**). Given that C62 and N34 can trigger the HNXX S-trimer to form post-fusion structures, albeit with markedly weaker triggering activity toward CV777, we hypothesized that these antibodies might function as artificial receptors when presented on the cell surface. To test this, we fused the TM of IL-2Rα to the C-terminus of the C62 or N34 heavy chain and co-expressed each membrane-anchored heavy chain with its corresponding light chain in HEK293T cells. In the presence of exogenously added trypsin, cell-surface-expressed C62 and N34 both markedly promoted CV777 S-protein-mediated cell-cell fusion, thereby mimicking the action of a natural receptor. In contrast, probably due to the HNXX S-protein already induced robust cell-cell fusion on its own, cell-surface expression of C62 or N34 did not further enhance fusion efficiency (**Fig. 5G**).

## Discussion

The neutralization mechanisms of antibodies against PEDV remain largely unexplored^18,20,21,42^. Based on studies of related coronaviruses, coronavirus-neutralizing antibodies generally act by blocking S-protein-mediated receptor binding^25,43–47^ or by interfering with S-protein-mediated membrane fusion^28,48–51^.

For PEDV, the functional receptor remains disputed^52–54^, which precludes direct assessment of whether neutralizing antibodies interfere with receptor binding. Here, we characterized the epitopes and binding modes of two S1^B^-targeting antibodies, C62 and N34. Although both antibodies showed highly compromised or no neutralizing activity against the G1 strain CV777, they differed markedly in their activity against the G2 strain HNXX: C62 effectively neutralized HNXX, whereas N34 showed little neutralizing activity (**Fig. 1A and C**). Because C62 and N34 bind distinct surfaces of S1^B^ (**Fig. 3D and H**), they could, in principle, interfere differently with receptor engagement if S1^B^ is confirmed to mediate receptor binding. However, in the absence of a confirmed PEDV receptor, this mechanism cannot be directly established.

Nevertheless, our data support a role for S-trimer premature triggering in C62-mediated neutralization^27^. C62 exhibited substantially stronger triggering activity than N34, robustly inducing proteinase K-resistant post-fusion S2 structures, whereas N34 showed much weaker activity (**Fig. 4A**). If premature S-trimer triggering is the principal mechanism of neutralization, insufficient triggering would be expected to compromise neutralizing potency, consistent with our observation with N34’s much weaker triggering activity (**Fig. 1A and 4**). Structural analysis indicates that, across multiple prefusion S-trimer conformations, the epitope and binding orientation of N34 would result in more extensive steric clashes than those of C62 (**Fig. 3C and I to K**). These clashes may restrict productive N34 binding onto the S-trimer and explain its weaker ability to trigger S-trimer fusion. Thus, the potent triggering activity of C62 likely underlies its effective neutralization of HNXX virus.

However, antibody neutralization mechanisms can be complex, particularly in the context of intact virions. Differences in S-trimer-binding mode and geometry may influence an antibody’s ability to cross-link S-trimers on the same virion or between different virions, thereby differentially affecting viral attachment, membrane fusion, and overall neutralization potency. This possibility is consistent with recent work showing that antibodies targeting distinct epitopes can differ in their ability to mediate S-trimer cross-linking and, consequently, in their effects on coronavirus neutralization^55^.

Interestingly, we also observed marked differences in infection phenotypes between the G2 strain HNXX and the G1 strain CV777. HNXX induced prominent syncytium formation, whereas CV777 did not (**Fig. 1A**). Consistently, HNXX S-protein showed robust activity in our cell-cell fusion assay, which measures cell-surface membrane fusion, whereas CV777 S-protein lacked detectable activity under the same conditions (**Fig. 5E and F**). Through domain-swapping experiments between HNXX and CV777 S-proteins, we found that determinants within the S1^A^ and S1^B^ cooperatively control PEDV S-protein-mediated cell-surface fusion activity (**Fig. 5E and F**).

The differential activity of C62 against HNXX and CV777 further suggests that the two S-proteins differ in their conformational dynamics. Although C62 binds more tightly to the isolated CV777 S1^B^ than to the HNXX S1^B^, it binds less efficiently to the intact CV777 S-trimer than to the HNXX S-trimer (**Fig. 1B**). This discrepancy suggests that the C62 epitope is less accessible in the context of the CV777 S-trimer, potentially because of differences in S1^B^ opening or overall spike dynamics. To explore the structural basis for this difference, we compared the S1^A^-S1^B^ interfaces in the HNXX and CV777 S-trimer structures in the 3-D0-up state, which has previously been associated with S1^B^ opening. The S1^A^-S1^B^ interface is markedly smaller in HNXX than in CV777, with buried surface areas of 23.9 Å² and 170.4 Å^2^, respectively (**Fig. S6**), indicating more extensive S1^A^-S1^B^ interactions in the CV777 spike. These differences may contribute to the distinct conformational dynamics of the two S-trimers and, consequently, to their different entry pathways and susceptibilities to antibody neutralization.

Although C62 showed much stronger activity than N34 in triggering HNXX S-trimer fusion in vitro, both antibodies functioned as artificial receptors in cell-cell fusion assays, promoting CV777 S-trimer-mediated fusion at the cell surface. This apparent discrepancy may reflect the experimental configuration of the cell-cell fusion assay, in which numerous surface-expressed S-trimer:antibody-artificial-receptor complexes are likely formed. Under these highly multivalent cell-cell engagement conditions, even the weak triggering activity of N34 may be sufficient to induce detectable cell-cell fusion. The ability of both antibodies to confer cell-surface fusion activity on the CV777 S-protein further suggests that ligands capable of altering S-protein conformational dynamics can modulate the spatiotemporal regulation of spike-mediated membrane fusion^27,56^.

Collectively, this study provides mechanistic insight into how two S1^B^-targeting PEDV antibodies, C62 and N34, differ in their neutralization activities. Our data also shed light on how determinants within the PEDV S-trimer govern cell-surface membrane fusion activity and may thereby shape PEDV entry pathway usage.

## Limitations and outlook

One important limitation is that the functional receptor for PEDV remains disputed. We were unable to detect binding between PEDV spikes and the disputed APN receptors, precluding direct assessment of whether C62 or N34 interferes with receptor engagement. While our data demonstrate that C62 neutralizes PEDV via premature triggering of S-trimers, we cannot formally exclude the possibility that C62 also contributes to neutralization by blocking receptor binding, should S1^B^ indeed function as the RBD. Despite this limitation, our structural and functional data support a model in which C62 neutralizes PEDV by prematurely triggering S-trimers. Our work also suggests that identifying antibodies capable of prematurely triggering S-trimers and promoting their disassembly may provide a generalizable strategy for discovering neutralizing antibodies against coronaviruses with unknown receptors.

## Materials and method

### Cell culture and virus preparation

Vero CCL81 and HEK293T cells were maintained in Dulbecco’s Modified Eagle Medium (DMEM; Gibco, USA) supplemented with 10% fetal bovine serum (FBS; OPCEL, China). Expi293F cells (Gibco, USA) were cultured in Expi293F Expression Medium (Gibco). All cells were incubated at 37 °C in a humidified atmosphere of 5% CO_2_, with Expi293F cultures being agitated in an orbital shaker. The PEDV strains CH/HNXX/2016 (GenBank accession No. MT338517.1) and CV777 (GenBank accession No. KT323979.1) were propagated and titrated in Vero cells.

### Virus neutralization assay

The neutralizing activity of the antibodies was evaluated using a modified neutralization assay as previously described^10^. Briefly, serial two-fold dilutions of the antibodies in DMEM were mixed with an equal volume of PEDV (strains HNXX or CV777) at a concentration of 4,000 TCID_50_/mL. The antibody-virus mixtures were incubated for 1 h at 37°C. Subsequently, 100 µL of each mixture was applied to confluent Vero cell monolayers in 96-well plates, followed by the addition of 100 µL of trypsin at a concentration of 20 µg/mL. Following a 48 h incubation at 37°C with 5% CO_2_, the cytopathic effect was observed under a microscope. The half-maximal inhibitory concentration (IC_50_) was determined by fitting the dose-response data to a four-parameter logistic regression model.

### Expression and purification of PEDV S-trimers and S1^B^

The codon-optimized extracellular domains of the S genes (from strains HNXX and CV777), fused with a T4 fibritin (foldon) trimerization domain and a C-terminal 6×His tag, were synthesized (BGI, China) and cloned into the pcDNA3.1 vector, generating plasmids pcDNA3.1-HNXX-S and pcDNA3.1-CV777-S. The S1^B^ gene fragments were subsequently subcloned from these plasmids into a pcDNA3.1 backbone containing a signal peptide and a 6×His tag, yielding plasmids pcDNA3.1-HNXX-S1^B^ and pcDNA3.1-CV777-S1^B^. All plasmids were transfected into Expi293F cells (density: 2.5×10^6^ cells/mL) using polyethyleneimine (PEI), and the cells were treated with kifunensine (final concentration of 5 μM) 3 h after transfection^14,15^. The culture supernatant was harvested 3–5 days post-transfection by centrifugation at 4,000 × g for 5 min and filtered through a 0.45 μm membrane. The filtrate was supplemented with 25 mM phosphate pH 8.0, 300 mM NaCl, 5 mM imidazole, and 0.5 mM PMSF, and then recirculated three times over a HiTrap TALON crude column (Cytiva, USA). The column was washed with 100 mL of buffer A (25 mM phosphate pH 8.0, 300 mM NaCl, 5 mM imidazole), before the target protein was eluted with a 100 mL linear gradient to 100% buffer B (25 mM phosphate pH 8.0, 300 mM NaCl, 500 mM imidazole). Target protein fractions were pooled, concentrated using an Amicon Ultra centrifugal filter [Merck Millipore; 10 kDa molecular weight cut-off (MWCO) for S1^B^ or 100 kDa MWCO for S-trimer], and buffer-exchanged into PBS.

### Antibody expression and purification

The variable heavy (VH) and variable light (VL) regions of monoclonal antibodies C62 and N34 (previously named as B34^9,10^) were cloned into pcDNA3.1 expression vectors containing the constant regions of porcine IgG1 heavy (GenBank: AK405771.1) and kappa light (GenBank: M59322.1) chains, respectively. For full-length IgG production, heavy-and light-chain plasmids were co-transfected at a 1:1 ratio into Expi293F cells using PEI, according to the manufacturer’s instructions. Antibodies were harvested from the culture supernatant five days post-transfection and purified using a Protein A affinity column (Cytiva). The purified IgGs were then concentrated and buffer-exchanged into phosphate-buffered saline (PBS) using a 50 kDa MWCO Amicon Ultra centrifugal filter (Merck Millipore). For Fab fragment generation, the CH2 and CH3 domains of the heavy chain were replaced with a C-terminal 6×His tag. The resulting Fab heavy-chain and light-chain plasmids were co-transfected (1:1 ratio) into Expi293F cells. The expressed Fabs were purified from the supernatant via Co^2+^ affinity chromatography (Cytiva), eluted with an imidazole gradient, and subsequently concentrated and buffer-exchanged into PBS using a 30 kDa MWCO Amicon Ultra device.

### Binding kinetics and affinity assessment by BLI

Binding kinetics and affinities of mAbs against the PEDV S-trimer or its S1^B^ subunit were quantified using BLI on an Octet R8 system (Sartorius). Competitive binding between S1^B^-targeting mAbs was assessed on the same platform, following an established protocol^28^. Briefly, Protein A biosensors were hydrated and equilibrated in kinetics buffer (PBS, pH 7.4, containing 1% BSA and 0.02% Tween-20). Biosensors were then loaded with IgGs at a concentration of 11 µg/mL. For affinity measurements, after loading IgG to a level of ∼1.5 nm, the association of analyte (S-trimer, S1^B^ subunit, or buffer control) was monitored for 300 s, followed by a 600 s dissociation phase in kinetics buffer. For competition assays, biosensors were first loaded with the S1^B^ subunit. A first mAb was then associated for 300 s, immediately followed by the association of a second mAb for another 300 s. All sensorgrams were processed using the FortéBio data analysis software HT v12.0.2.59 (Sartorius) to determine the dissociation constants (*K*_D_).

### Negative-stain electron microscopy

Purified PEDV HNXX S-trimer proteins (1 mg/mL) were incubated with either IgGs (C62, N34) at a 1:3 molar ratio, 1 h at 25°C. Samples were diluted to a final S-protein concentration of 0.05 mg/mL, and 5 µL aliquots were applied to glow-discharged (15 mA, 45 s), carbon-coated copper grids and allowed to adsorb for 1 min. Excess liquid was blotted with filter paper, and grids were stained twice with 0.75% (w/v) uranyl formate. Micrographs were acquired on a JEM-1400 FLASH transmission electron microscope (JEOL, Japan) operated at 120 kV and equipped with a 2560×1920 XAROSA 20 megapixel CMOS camera (Emsis, Germany), using RADIUS imaging and analysis software at a nominal magnification of ×50,000.

### Western blot

Three aliquots of purified PEDV S-trimer (50 nM, based on S-protomer molecular weight) from strains HNXX or CV777 were incubated with either 50 nM of monoclonal antibodies (C62 or N34) or an equal volume of PBS (negative control, pH 7.4) for 1 h at 25°C. After incubation, one aliquot from each treatment group was immediately mixed with SDS-PAGE loading buffer and heat-denatured (98°C, 10 min) to serve as the undigested control. The remaining two aliquots were subjected to trypsin digestion (50 µg/mL, 37°C, 1 h) in PBS. Following trypsin treatment, one of the two digested aliquots was further digested with Proteinase K (50 µg/mL, 37°C, 1 h) in the same buffer. All enzymatic reactions were terminated by the addition of loading buffer and immediate heat denaturation at 98°C for 10 min. Processed protein samples were separated by SDS-PAGE on 4–12% gradient gels and transferred onto PVDF membranes. For immunodetection, membranes were probed with rabbit polyclonal anti-PEDV S2 antibodies (PVV31306, AntibodySystem, France), followed by HRP-conjugated goat anti-rabbit IgG secondary antibody. Protein signals were visualized using enhanced chemiluminescence.

### IFA

To evaluate the effects of camostat and E-64d on viral infection, Vero cells were pretreated with each compound for 1 h prior to viral infection. At 12 h post-infection (hpi), the cells were washed three times with PBS and fixed with 4% paraformaldehyde (Biosharp, China) for 30 minutes at room temperature. After three additional PBS washes, the cells were permeabilized with 0.2% Triton X-100 for 10 minutes at room temperature. Subsequently, the cells were incubated with an anti-PEDV N protein antibody (Qianxun Biotech, China; 10 μg/mL) for 1 hour at 37°C, followed by a 30 min incubation with a Cy3-conjugated goat anti-mouse IgG secondary antibody (Abclonal, China; 1:200 dilution) at 37°C. Following a washing step, the nuclei were stained with DAPI for 10 minutes at room temperature. After a final wash, the samples were imaged using a fluorescence microscope.

### qPCR

To assess the antiviral activity of camostat and E64d, Vero cells were pretreated with each compound for 1 hour before infection with PEDV. Cells were harvested at 12 h post-infection (hpi), and total RNA was extracted for analysis. The relative mRNA levels of the PEDV S gene were quantified by qPCR, normalized to GAPDH, and calculated using the 2^−ΔΔCT^ method. Reverse transcription was performed with a commercial kit (Abclonal) under the following conditions: 37°C for 2 min, 55°C for 15 min, and 85°C for 5 min. The subsequent qPCR protocol comprised an initial denaturation at 95°C for 10 min, followed by 40 cycles of 95°C for 5 s and 60°C for 30 s.

### Assessment of S-protein expression by flow cytometry

Cell surface expression of the S-protein in the membrane fusion assay was quantified by flow cytometry. Briefly, cells were washed three times with washing buffer (PBS containing 1% BSA) and then incubated with an anti-PEDV S2 antibody^21^ (20 µg/mL) for 2 h on ice. Following primary antibody incubation, the cells were washed and stained with Cy3 or FITC-conjugated goat anti-mouse IgG secondary antibody (Abclonal; 1:200 dilution) for 1 h on ice. After a final wash, the cells were resuspended in washing buffer and analyzed on a CytoFLEX LX flow cytometer (Beckman Coulter, USA). A minimum of 10,000 events were acquired for each sample, and data analysis was performed using FlowJo VX software.

### Sequences alignments

The phylogenetic analysis was conducted using 95 S gene sequences of various PEDV strains retrieved from GenBank (Table S2), with the Maximum Likelihood method implemented in MEGA software. To assess the conservation of the N34 and C62 antigenic epitopes, eleven representative amino acid sequences spanning distinct phylogenetic subgroups were aligned and analyzed using ESPript (https://espript.ibcp.fr/ESPript/ESPript/).

### Cell-cell fusion assay

The complementary split-GFP fragments, GFP1–10 and GFP11, were expressed in effector and target cells, respectively, by transfection, enabling reconstitution of functional GFP upon cell-cell fusion^57^. HEK293T cells expressing the S-proteins and GFP1–10 were prepared as effector cells and HEK293T cells expressing the GFP11 without overexpressing any receptor proteins were prepared as target cells. Briefly, HEK293T cells were cultured in six-well plates to 70–80% confluence and transfected with 2 μg of pQCXIP-GFP1-10 and 2 μg of pcDNA3.1(+)-HNXX-S using 8 μL of jetPRIME reagent to generate effector cells. In parallel, HEK293T cells were transfected with 2 μg of pQCXIP-BSR-GFP11 using 4 μL of jetPRIME reagent to generate target cells. For the artificial receptor assay, the TM of IL-2Rα was fused to the C-terminus of the C62 or N34 heavy chain^56^. These membrane-anchored heavy chains were co-expressed with their corresponding light chains and GFP11 in HEK293T cells, which were used as target cells. After 24 h, both effector and target cells were washed, resuspended in DMEM, mixed at a 1:1 ratio, and seeded at 2 × 10^5^ cells per well in 96-well plates. Fluorescence images were acquired at the indicated time points using a fluorescence microscope. For antibody inhibition assays of membrane fusion, effector cells expressing the S-protein were pre-incubated with the indicated antibody at 37 °C for 1 h before co-culture with target cells.

### Cryo-EM sample preparation and data collection

For the C62 Fab:S1^B^:N34 Fab ternary complex, protein components were mixed at 1:1:1 molar ratio for 1 min, before 3 µL of the mixture was applied to glow-discharged (15 mA, 30 s) Quantifoil Au R1.2/1.3 holey carbon grids. All grids were blotted for 3 s (force 4) and plunge-frozen in liquid ethane using a Vitrobot Mark IV (Thermo Fisher, USA) at 4°C and 100% humidity. The grids were imaged on a 300 kV Titan Krios (Thermo Fisher) with Falcon4 detector and SelectrisX energy filter (10 eV slit width). Data were collected in EER mode at ×165,000 magnification (0.73 Å pixel size) using EPU, with defocus –0.8 to –2.4 µm. Gain-normalized movies (30 frames, ∼50 e⁻ Å⁻^2^ total dose) were recorded in counting mode.

### Cryo-EM data processing

The C62 Fab:S1^B^:N34 Fab ternary complex dataset was processed as follows: Movie frames were motion-corrected and dose-weighted, followed by CTF estimation^58,59^. An initial particle set (from 500 micrographs) was identified using reference-free blob picking. After particle extraction and 2D classification, well-resolved class averages served as templates for subsequent particle picking. Selected particles underwent ab initio reconstruction and 3D classification. The highest-quality 3D class was used to train Topaz for additional particle picking. Following three iterative rounds of particle picking and 3D classification, we obtained an initial reconstruction at 3.66 Å resolution. Final refinement incorporated orientation diagnostics, 3D flexibility analysis, and non-uniform refinement, yielding a final map at 3.0 Å resolution. Local resolution variations were assessed using the 3D-FSC method.

### Model building and analysis

AlphaFold Server (https://alphafoldserver.com) was used to generate the S1^B^ and Fab models using the amino acid sequences of the PEDV HNXX S-protein and the heavy and light chains of the C62 and N34 Fabs, respectively. The structures created by AlphaFold was fitted into the C62 Fab:S1^B^:N34 Fab map. The structure was manually adjusted in Coot 0.9.6^60^ and refined in PHENIX 1.20.1^61^. Real-space refinement was carried out iteratively in Coot and PHENIX. Model refinement statistics are summarized in Table S3. Interfaces analysis was performed by PISA^62^. Figures were generated in UCSF Chimera^63,64^.

## Data availability

Cryo-EM density map for C62 Fab:S1^B^:N34 Fab have been deposited in the Electron Microscopy Data Bank (EMDB) with accession codes EMD-67251. Related atomic model has been deposited in the Protein Data Bank (PDB) under the accession code 9XTU, respectively. Source data are provided with this paper.

## Acknowledgements

The research is supported by the Prevention and Control of Emerging and Major Infectious Diseases-National Science and Technology Major Project (2026ZD01999601 to X.X.); National Natural Science Foundation of China (82341085 and 32570199 to X.X.); the National Key R&D Program of China (2021YFA1300903 to X.X.); Major Project of Guangzhou National Laboratory (GZNL2024A01010 to X.C.); Major Project of Guangzhou National Laboratory (SRPG22-002 to X.X.); Basic Research Project of Guangzhou Institutes of Biomedicine and Health, Chinese Academy of Sciences (GIBHBRP24-02 to X.X.); Science and Technology Planning Project of Guangdong Province, China (2023B1212060050 to X.X.). X.X. acknowledges Start-up grants from the Chinese Academy of Sciences; the Fundamental Research Fund for the Central Universities (2662025DKPY009) and the earmarked fund for CARS-41 to H.Z.

## Author Contributions

J.L., S.W., X.C., H.Z., and X.X. conceived and designed the study; J.L., and S.W. performed the experiments; J.L., S.W., Z.L., and X.X. determined cryo-EM structures; J.L., S.W., and X.X prepared the figures; S.W., Z.W., Y.L., J.Z., and H.Z. prepared the antibodies. S.W., J.L., and X.Z. performed the virus neutralization experiments; S.X., Y.M., and H.Z. contributed materials; J.L., S.W., and X.X. wrote the initial manuscript, which was revised by J.L., S.W., and X.X. with contributions from all authors.

---

**Fig. S1.**
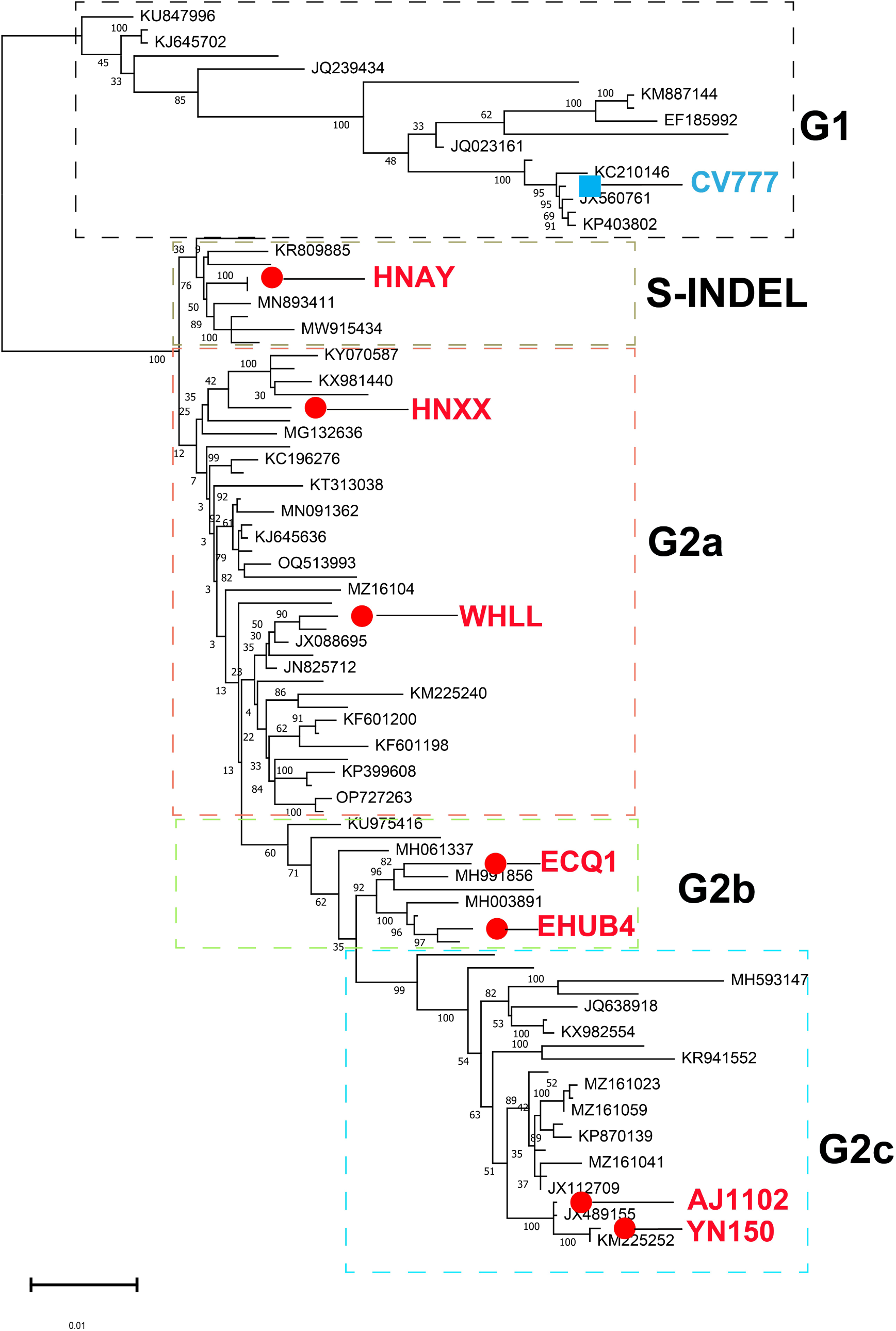
Phylogenetic analysis of PEDV S genes. The maximum-likelihood analysis was performed using MEGA with the default gap-opening penalty of 15. The 95 PEDV S-gene sequences included in the analysis are summarized in Table S2. Red dots indicate G2 strains tested in previous neutralization assays, and the blue square indicates the tested G1 strain.

**Fig. S2.**
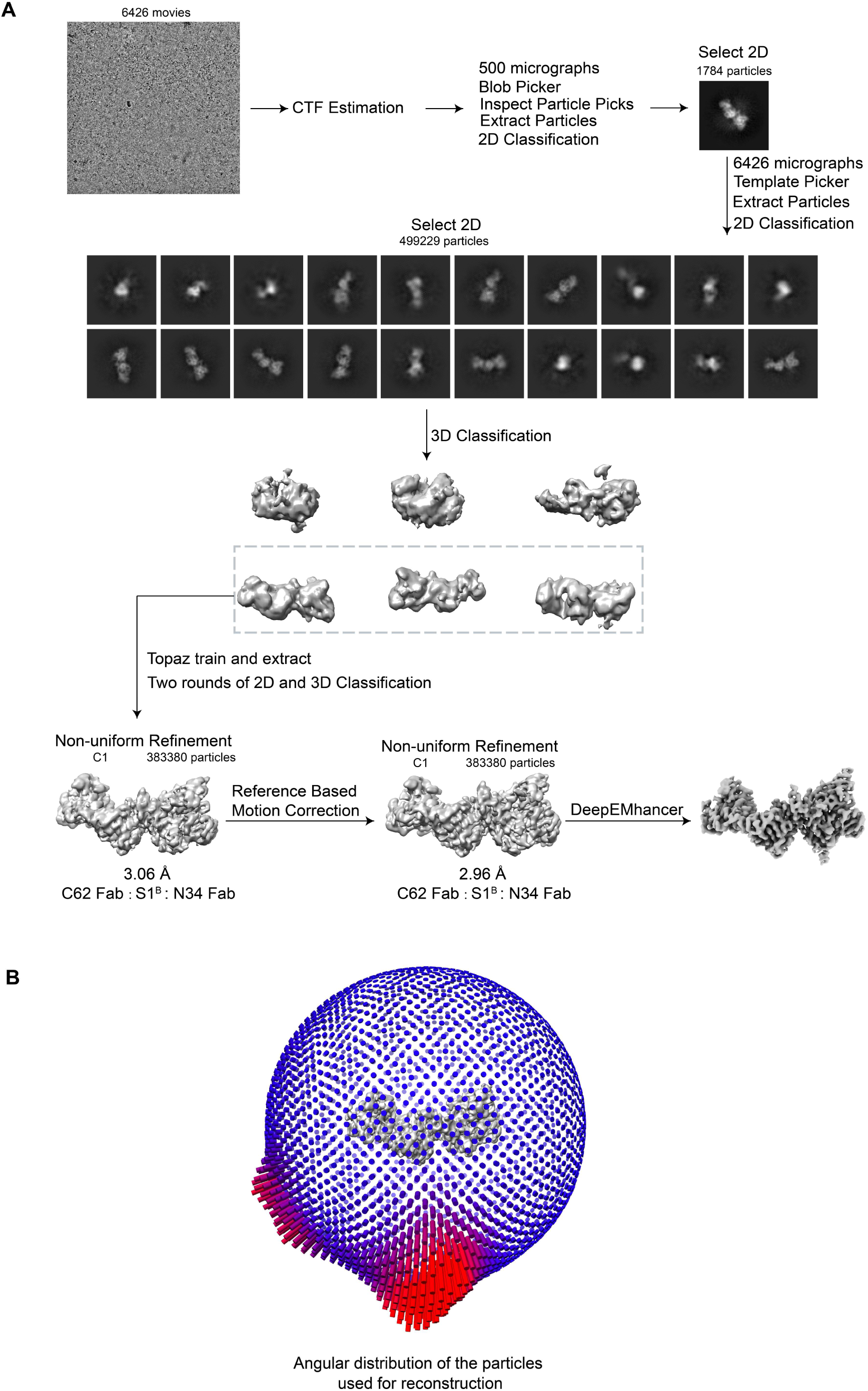
Cryo-EM data processing flow-chart for the C62 Fab:S1^B^:N34 Fab complex structure. **A,** Following initial particle selection through blob picking, multiple rounds of 2D and 3D classification were conducted to remove contaminating particles. The selected particles were reconstructed using Non-uniform Refinement. After reference based motion correction was performed on the resulting map, the particles were finally reconstructed again using Non-uniform Refinement. **B**, Angular distribution of the particles used for the reconstruction of the C62 Fab:S1^B^:N34 Fab complex structure.

**Fig. S3.**
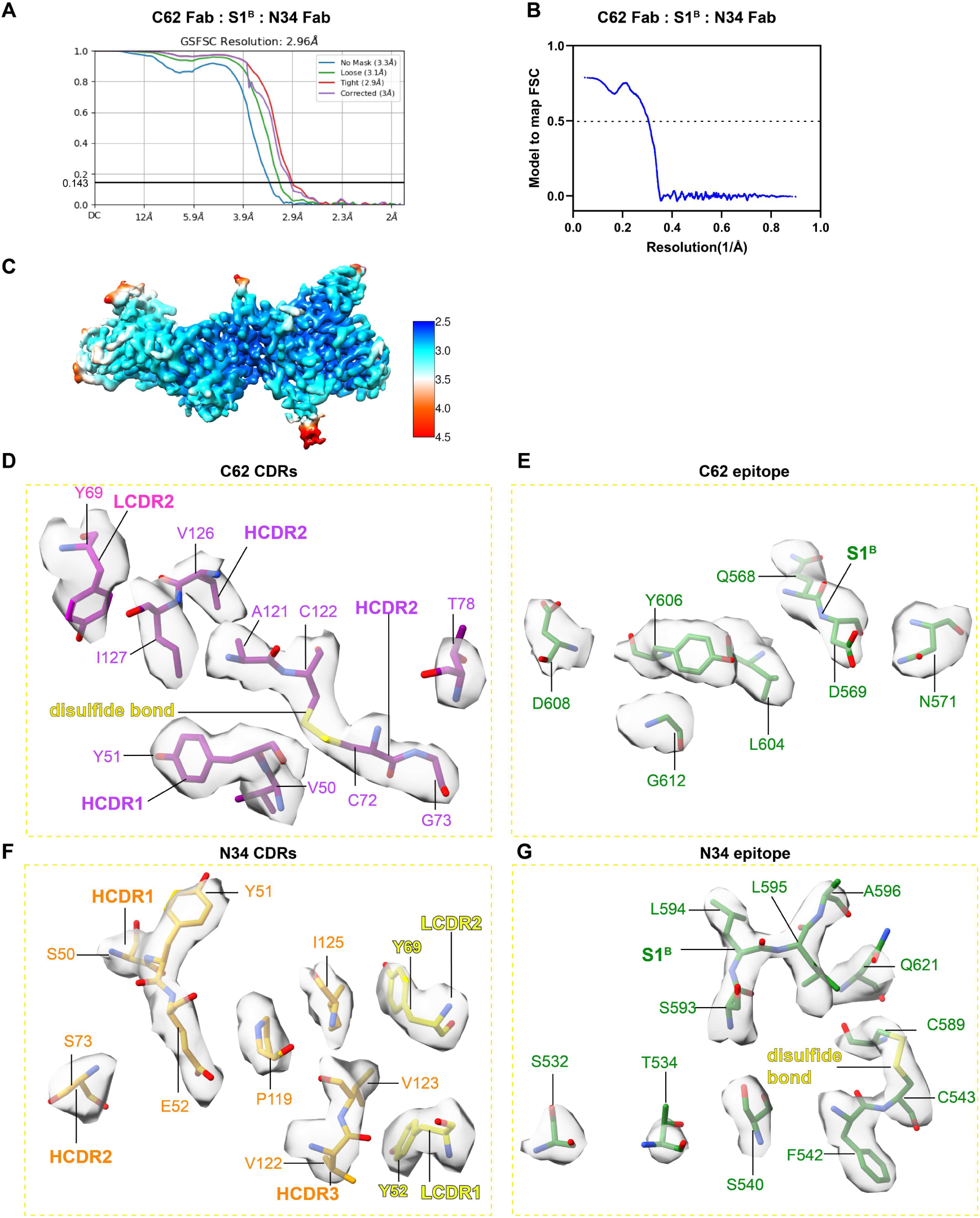
Resolution assessment of the cryo-EM structure. **A**, Global resolution assessment by Fourier shell correlation (FSC) at the 0.143 criterion. **B**, Correlations of model vs map by FSC at the 0.5 criterion. **C**, Local resolution map for the C62 Fab:S1^B^:N34 Fab complex. **D** and **E**, Representative densities of the C62 epitope and C62 CDRs loops in the final map. C62 LCDRs, HCDRs, and epitopes are colored in magenta, purple and green, respectively. **F** and **G**, Representative densities of the N34 epitope and N34 CDRs loops in the final map. N34 LCDRs, HCDRs, and epitopes are colored in yellow, brown and green, respectively.

**Fig. S4.**
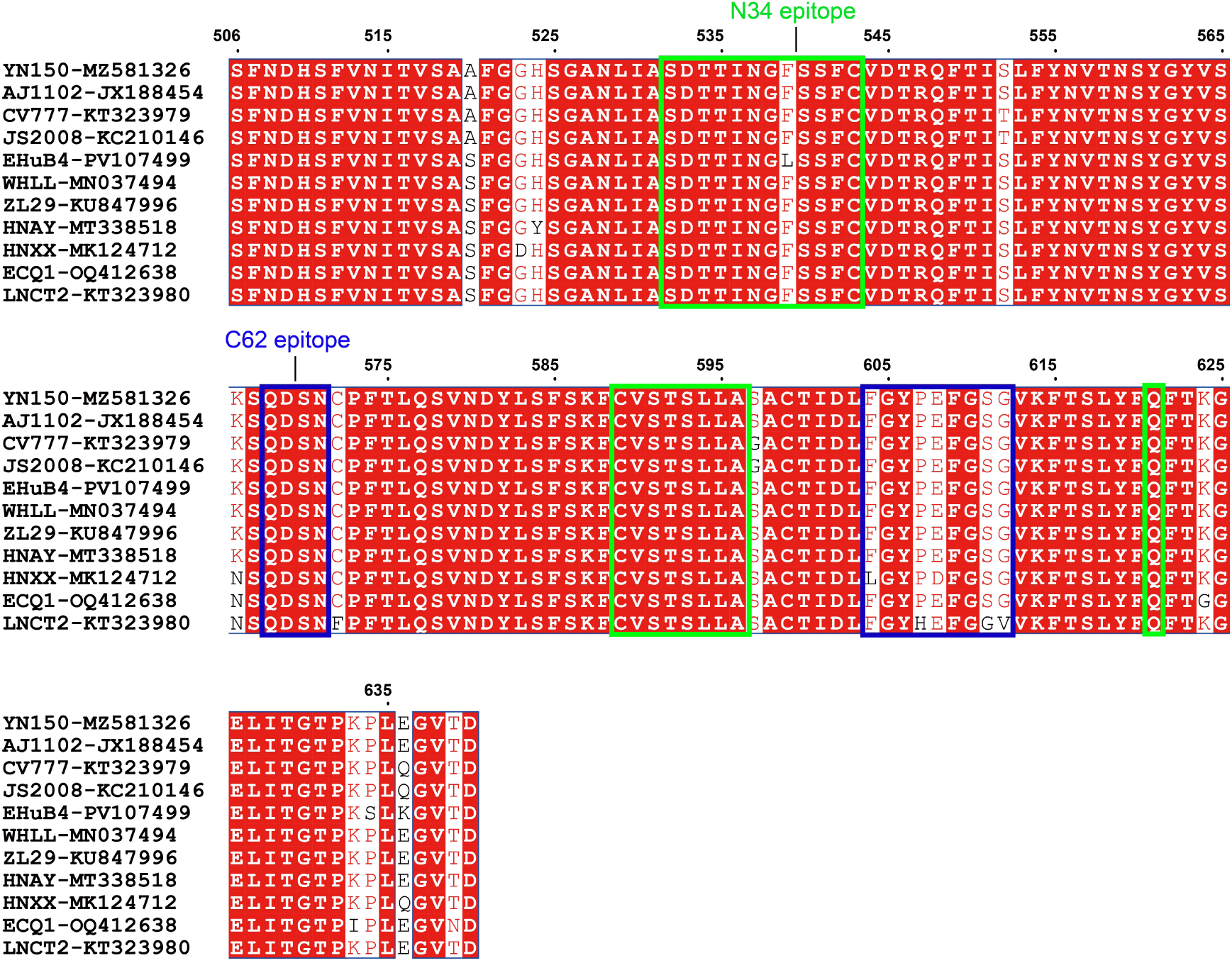
Conservation analysis of antigenic epitopes. Multiple sequence alignment of S1^B^ sequences from PEDV strains YN150, HNXX, WHLL, AJ1102, EHuB4, HNAY, ECQ1, ZL29, LNCT2, CV777, and JS2008. Amino acid residues within the two blue boxes constitute the C62 epitope. Amino acid residues within the three green boxes constitute the N34 epitope.

**Fig. S5.**
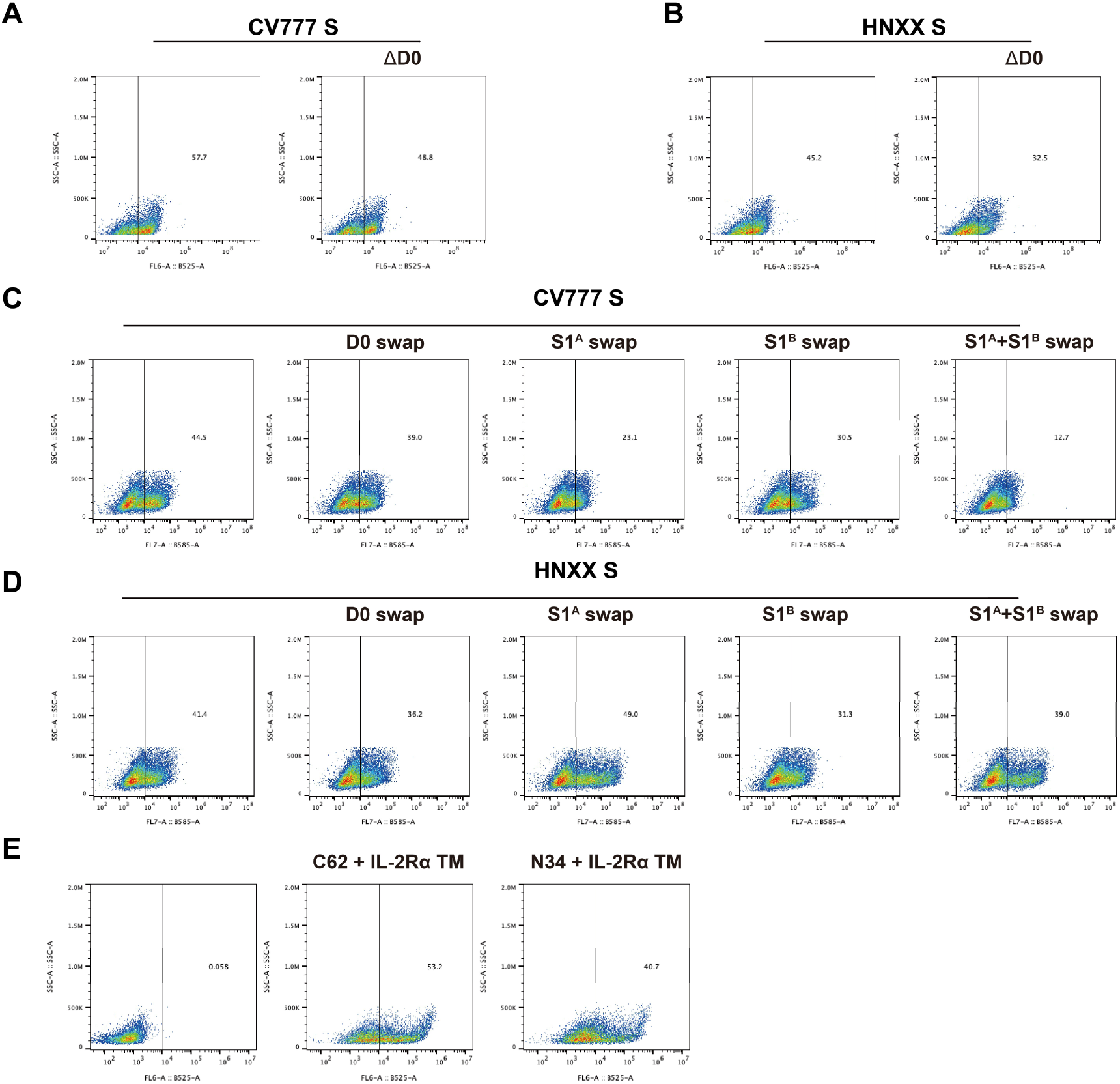
Cell surface spike expression by flow cytometry. **A**, HEK293T cells were transfected with either pcDNA3.1-CV777-S or pcDNA3.1-CV777-S-ΔD0 (D0-deletion) plasmids. **B**, HEK293T cells were transfected with either pcDNA3.1-HNXX-S or pcDNA3.1-HNXX-S-ΔD0 plasmids. Expression of all proteins was confirmed by flow cytometry. **C** and **D**, HEK293T cells were transfected with the domain-swap plasmids of CV777 and HNXX S-proteins (illustrated in Fig. 5D), involving swapping of the D0, S1^A^, S1^B^, or S1^A^+S1^B^ regions, respectively. **E**, HEK293T cells were transfected with plasmids engineered to express C62 IgG or N34 IgG with the IL-2Rα TM domain fused to the C-terminus of their heavy chains.

**Fig. S6.**
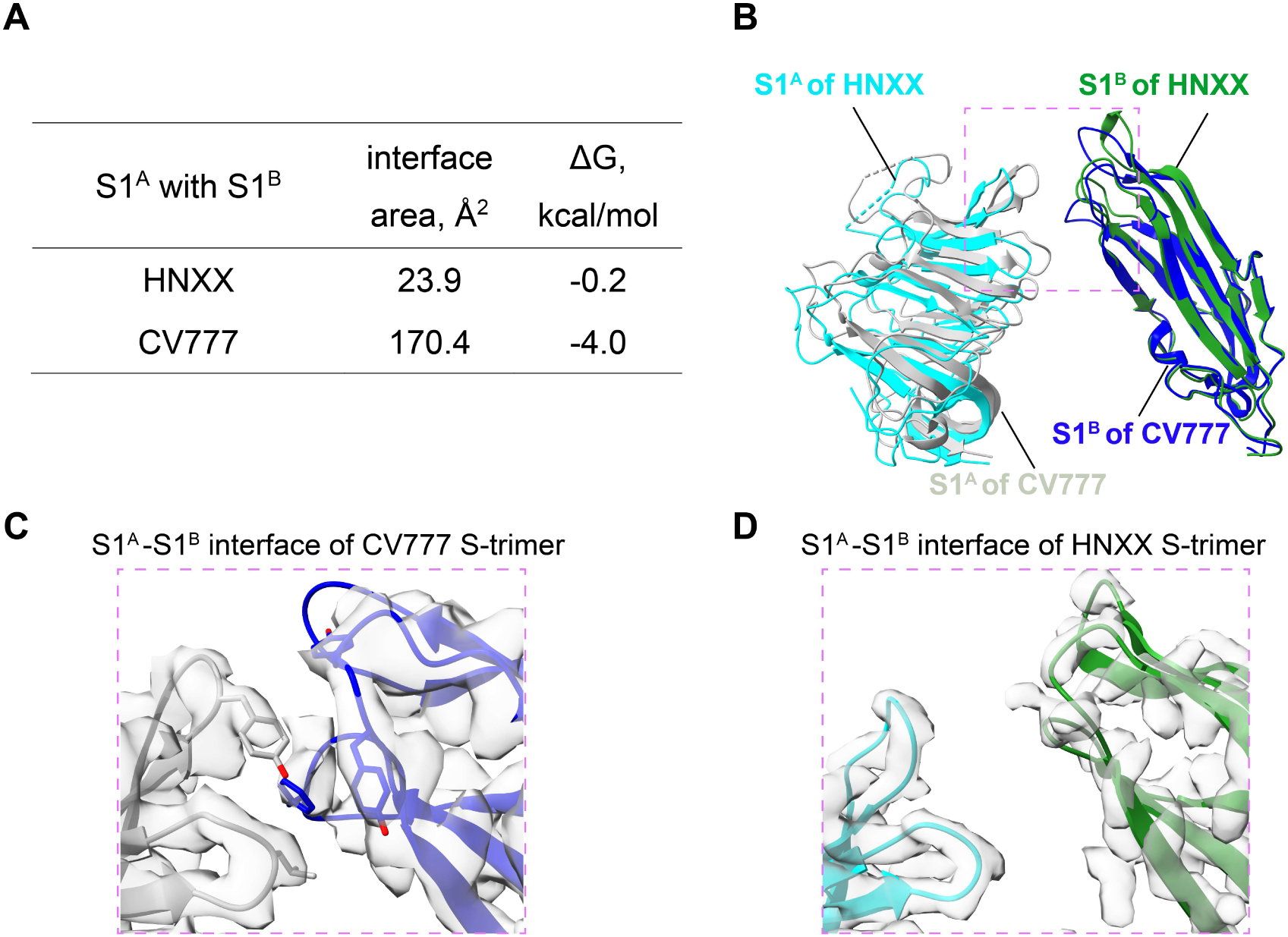
S1^A^-S1^B^ interface of the HNXX and CV777 spike proteins. **A**, Interface areas and solvation free energies (ΔG) of the S1^A^-S1^B^ interfaces in the HNXX (S-trimer in D0_UUU_, PDB: 9XNZ) and CV777 (S-trimer in D0_UUU_, PDB: 6U7K) S-trimers, calculated using PDBePISA. **B**, Structural alignment of the S1^A^-S1^B^ interfaces, using HNXX S1^B^ (D0_U_) as the reference. HNXX S1^A^ (cyan) and S1^B^ (dark green), and CV777 S1^A^ (gray) and S1^B^ (blue) are shown as cartoons. **C** and **D**, Representative cryo-EM densities at the S1^A^-S1^B^ interfaces are shown for the CV777 S-trimer and HNXX S-trimer, respectively.

**Table S1.**
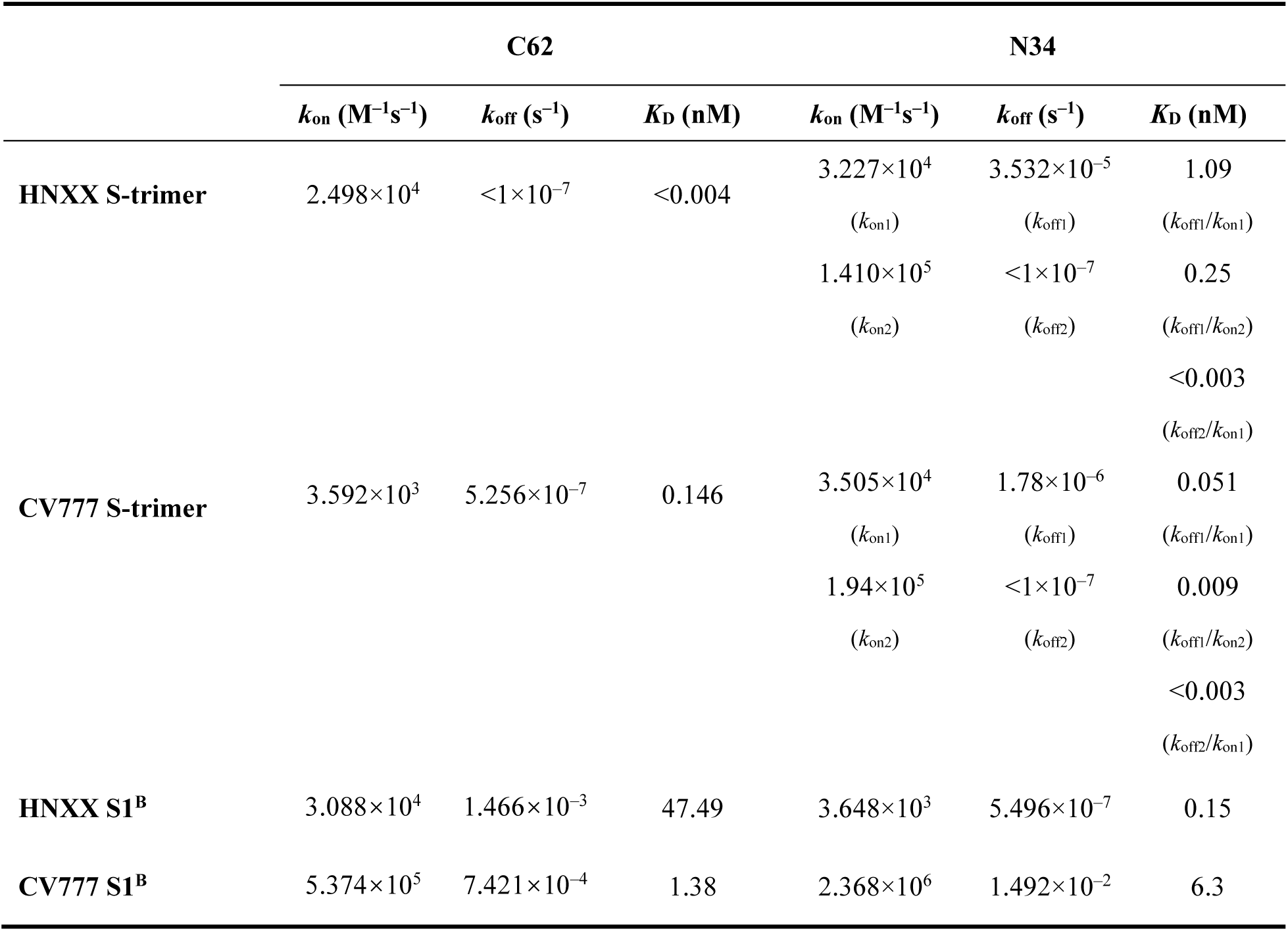
Kinetic parameters for antibody binding to PEDV S trimers and S1^B^, as determined by BLI.

**Table S2.** The S genes of PEDV strains used for phylogenetic analysis.

| GenBank No. |  |  |
| --- | --- | --- |
| MN091362.1 | MZ581326.1 | KM225244.1 |
| KT313038.1 | JX188454.1 | KF601198.1 |
| KY211062.1 | OQ412638.1 | MK507906.1 |
| MH991864.1 | PV107499.1 | OP727263.1 |
| MK532989.1 | MT338518.1 | OQ349215.1 |
| MK111633.1 | MT338517.1 | OQ513993.1 |
| MN893411.1 | MN037494.1 | KF601200.1 |
| KM406180.1 | AB548618.1 | KF601201.1 |
| MW915434.1 | KM887144.1 | KX073610.1 |
| MK533010.1 | JN547228.1 | KP399608.1 |
| KY007139.1 | GU937797.1 | KP399630.1 |
| KY007140.1 | KT323979.1 | MZ16104.1 |
| PX842723.1 | JQ023161.1 | KJ646582.1 |
| PP461398.1 | AB857235.1 | KM225240.1 |
| KT323980.1 | KP403802.1 | KX982554.1 |
| MF038016.1 | KC210146.1 | KR941552.1 |
| MW915432.1 | KF898124.1 | OK642747.1 |
| OP235511.1 | EF185992.1 | KM225252.1 |
| KF601199.1 | JX560761.1 | KF468753.1 |
| MH003891.1 | KU646831.1 | MZ161024.1 |
| MK532999.1 | JN825712.1 | MZ161059.1 |
| KX982576.1 | KJ526096.1 | MZ161041.1 |
| MH991856.1 | KC196276.1 | KU975416.1 |
| KY619768.1 | KR809885.1 | MK685665.1 |
| KP870139.1 | KX981440.1 | KJ399978.1 |
| MZ161023.1 | MH053413.1 | KJ645702.1 |
| JQ638918.1 | JX261936.1 | KU847996.1 |
| KR265796.1 | MH593147.1 | JX112709.1 |
| KJ645636.1 | MH061337.1 | JX088695.1 |
| KP870113.1 | KP765609.1 | KY070587.1 |
| JQ239434.1 | JX647847.1 | JX489155.1 |
| KP276248.1 | MG132636.1 |  |

**Table S3.** Cryo-EM data collection, refinement and validation statistics.

| C62 Fab:S1B:N34 Fab<br>(EMD-67251, PDB: 9XTU) |  |
| --- | --- |
| **Data collection and processing** |  |
| Magnification | 130000 |
| Voltage (kV) | 300 |
| Electron exposure (e-/Å<sup>2</sup>) | 50 |
| Defocus range (μm) | −0.8 to −2.4 |
| Pixel size (Å) | 0.93 |
| Movies (no.) | 6433 |
| Initial particle images (no.) | 499229 |
| Symmetry imposed | C1 |
| Final particle images (no.) | 383380 |
| Map resolution (Å) | 3.0 |
| FSC threshold | 0.143 |
| Map resolution range (Å) | 2.6 to 42.2 |
| **Refinement** |  |
| FSC threshold | 0.5 |
| Model resolution (Å) | 3.28 |
| Map sharpening *B* factor (Å<sup>2</sup>) | −139.3 |
| **Model composition** |  |
| Non-hydrogen atoms | 5161 |
| Protein residues | 685 |
| Ligands | 2 |
| B factors (Å<sup>2</sup>) |  |
| Protein | 27.31 |
| Ligand | 24.64 |
| **R.m.s. deviations** |  |
| Bond lengths (Å) | 0.004 |
| Bond angles (°) | 0.776 |
| **Validation** |  |
| MolProbity score | 1.75 |
| Clashscore | 5.63 |
| Poor rotamers (%) | 0.00 |
| **Ramachandran plot** |  |
| Favored (%) | 93.02 |
| Allowed (%) | 6.98 |
| Disallowed (%) | 0.00 |

